# Attenuation of Inflammatory Cytokines by Selective Vagal Motor Stimulation via Silicone Spiral Nerve Cuff

**DOI:** 10.64898/2026.06.10.731452

**Authors:** Kathryn S. Mirandette, Atharva Sahasrabudhe, Magdalena Slowikowski, John H Caldwell, Polina Anikeeva, Richard F. ff. Weir, Arjun K. Fontaine

## Abstract

**Objectives:** To determine how selective optogenetic vagus nerve stimulation (VNS) of distinct axonal subpopulations modulates systemic inflammatory cytokines in an acute model of endotoxemia.

**Materials and Methods:** A silicone spiral nerve cuff with integrated custom probes including microscale light-emitting diodes (μLEDs) was fabricated and implanted on the left cervical vagus nerve of anesthetized transgenic mice expressing ChR2 under cholinergic (ChAT) or glutamatergic (Vglut2) cell promoters. Lipopolysaccharide (3 mg/kg) was administered intraperitoneally to induce endotoxemia, and mice received optical VNS for 2 hours. Blood was collected 30 minutes after VNS termination and quantified via immunoassay for serum inflammatory cytokines (IL-6, IL-1β, TNF-α, IL-10) and C-reactive protein (CRP).

**Results:** ChAT-selective optical VNS significantly reduced IL-6 (p = 0.027) and IL-1β (p = 0.026) relative to Cre-negative sham controls. Vglut2-targeted stimulation did not significantly reduce IL-6, IL-1β, or TNF-α versus sham. Cytokine levels were significantly reduced with ChAT VNS compared to Vglut2 VNS in all pro-inflammatory cytokines (IL-6: p=0.024, IL-1β: p=0.011, TNF-α: p=0.030). The anti-inflammatory cytokine IL-10 was significantly decreased with ChAT versus Vglut2 VNS (p=0.033). CRP levels were not statistically different between groups.

**Conclusions:** Optogenetic VNS targeting cholinergic neurons produced robust suppression of key pro-inflammatory cytokines IL-1β and IL-6, whereas stimulation of glutamatergic neurons did not significantly alter inflammatory cytokine levels, highlighting the importance of pathway selectivity in the inflammatory effects of VNS. These findings highlight cell-type specific optogenetic neuromodulation as a valuable tool for assessing impact of vagal circuits and support preferential targeting of efferent cholinergic neurons in acute systemic inflammation.

## Introduction

The vagus nerve is the main parasympathetic branch of the autonomic nervous system^1^ with extensive innervation throughout the visceral organs^2^. Vagal fibers are diverse, ranging from unmyelinated fibers < 0.5 μm in diameter to myelinated fibers > 10 μm in diameter.^3^, Given its extensive innervation, the vagus nerve represents a promising target for neuromodulation therapies. Vagus nerve stimulation (VNS) is approved by the U.S. Food and Drug Administration (FDA) for drug-resistant epilepsy^4^, treatment-resistant depression^5^, Rheumatoid Arthritis (RA)^6^ and stroke rehabilitation^7^, while many clinical trials are exploring the efficacy of VNS in treating conditions such as migraines^8^, Crohn’s disease^9, 10^, and obesity^11^. The vagus nerve’s influence on immune signaling, specifically, the regulation of inflammatory mediators, has been well established.^1^ Chronic inflammation is increasingly recognized as an underlying factor in a wide range of diseases including autoimmune conditions (e.g. RA, Crohn’s disease, multiple sclerosis), neurodegenerative diseases such as Alzheimer’s, various cancers, diabetes, and severe, life-threatening conditions such as sepsis.^12, 13^ Chronic inflammatory conditions contribute to ∼ 60% of deaths globally.^14^

Inflammatory processes are regulated, in part, by the vagus nerve. Sensory pathways of the vagus nerve detect inflammatory stimuli, including cytokines and other pathogen derived products, and convey these signals to the brain within milliseconds.^12^ Outgoing (efferent) neural pathways can, in turn, modulate inflammatory signaling in the body; the cholinergic anti-inflammatory pathway (CAP) of the vagus nerve down-regulates the production of systemic proinflammatory cytokines^15, 16^ and modulates local tissue inflammation.^17, 18^ Given these inflammation modulating pathways, the vagus nerve provides an important target to balance dysregulated inflammation. Prior work suggests that efferent and afferent vagus nerve stimulation recruit distinct neuroimmune pathways.^19, 20^ Activation of afferent vagal fibers can attenuate immune responses through downstream pathways^21^, including reflex activation of cholinergic fibers as well as spinal sympathetic circuits^22^, whereas stimulation of vagal efferent fibers can directly engage effector mechanisms, such as the CAP, to effect inflammatory output.^15, 23^ In RA patients, VNS inhibits production of TNFα, IL-1β, and IL-6 and improves symptoms^24^, while VNS exerts anti-inflammatory impact^15, 16, 25–30^ in patients with Crohn’s disease^31^, in models of pancreatitis^32^, inflammatory bowel disease (IBD)^30, 33^, and acute lung inflammation^19, 24, 31^.

To modulate nerve activity, electrode-based systems that interface with neural tissue are commonly employed. However, a major limitation of electrical stimulation is its lack of cell-type specificity. All neurons, regardless of cell type or function, within the stimulation field exceeding the depolarization threshold are activated. This lack of specificity can lead to off-target activation, which can result in adverse effects including cough, hoarseness, voice alteration, throat and neck pain, headache, and dyspnea.^34^ In addition, activation of aggregate pathways precludes the ability to discern and understand the impactful circuits which are most therapeutically relevant. An alternate method of interfacing with neural tissue is via optogenetics. Optogenetic techniques offer genetically-defined cell-type specificity and millisecond-scale temporal control over neural circuit activation or inhibition, providing a level of precision unattainable with traditional electrical stimulation.^35, 36^

Because the precise mechanisms of VNS that alleviate clinical symptoms are often not well understood, pathway-selective targeting within the vagus nerve is needed to better characterize therapeutically impactful axonal subsets to optimize clinical benefit and deliver treatments without off-target effects. In the current study, we investigated how cell type-specific stimulation of efferent and afferent vagal fibers in a model of systemic inflammation modulates key inflammatory cytokines. We designed and fabricated a silicone nerve cuff with embedded microscale light-emitting diodes (µLEDs) that provides a biocompatible interface with the mouse vagus nerve, enabling precise optogenetic activation of defined vagal fiber subsets.

## Materials and Methods

### Silicone Spiral Cuff Fabrication

The cuff design and fabrication method was inspired by work from Naples et al.^37^, adapted to fit our design requirements and specifications. Silicone cuffs were fabricated with Silastic® MDX4-4210 BioMedical Grade Elastomer (MDX) (Dow Inc.). The two-part silicone system was mixed in a 10:1 base:curing-agent ratio by weight according to the manufacturer’s protocol. To remove air bubbles, the mixture was centrifuged at 2000 x g for 30 seconds and placed under vacuum at 25 psi for 45 minutes. The silicone spiral was generated using two layers of silicone. The first sheet was molded between two 2 × 3-inch glass slides separated by stainless steel shim stock (76.5 μm). A thin layer of mold release (EASE RELEASE™ 205; Mann Release Technologies Inc) was applied to the interior of the mold to facilitate demolding. The MDX was cured at 100°C for 20 minutes and demolded, yielding a 2 × 2-inch silicone base sheet. The base sheet of cured MDX was clamped in a strain device and held at 250% percent strain under uniform tension. A second layer of prepared MDX was poured over a 1 × 1 inch area on the stretched base sheet between spacers (76.5 μm). This layer was cured at room temperature for 24 hours under a weighted glass slide coated in mold release. The resulting spiral sheet was trimmed to approximately 1-2mm in width (*Supplementary Figure S1*).

### Micro-LED Probe Fabrication & Integration

A glass slide was used as the carrier substrate after sequential cleaning with acetone, isopropyl alcohol (IPA), and UV-ozone treatment. A thin sacrificial layer of poly(vinyl alcohol) (PVA, Sigma) was then spin-coated onto the glass and baked on a hot plate at 120 °C for 15 min. Next, an approximately 100 µm-thick layer of polydimethylsiloxane (PDMS, 10:1 base:curing agent) was spin-coated and cured at 90 °C for 1 h. A copper-clad polyimide sheet, consisting of 9 µm Cu on 25 µm polyimide, was UV-ozone treated, laminated onto the PDMS layer, and baked at 120 °C for 30 min to improve adhesion. The Cu layer was then selectively patterned into the desired layouts, followed by low-temperature solder bonding of µLEDs (Cree Inc., TR2227, 475 nm emission) using low-temperature ChipQuik solder paste. A 14 to 20 µm-thick parylene-C layer was subsequently deposited by chemical vapor deposition to serve as a fluid and moisture barrier for the underlying electronics. An additional PDMS layer, approximately 50 µm thick, was then coated and cured at 90 °C for 1 h. Electrical access pads were opened by selective laser ablation through the insulating layers. Device outlines were then defined by laser ablation, followed by release of the devices from the glass carrier by dissolution of the sacrificial layer in deionized water at 120 °C for 12 h. Finally, insulated stainless-steel wires were soldered to the Cu access pads, and the interconnect regions were encapsulated with UV-curable epoxy (Norland) to complete device fabrication.

The µLED probe was integrated within the inner diameter of the silicone spiral cuff using additional MDX, cured at 50 °C for three hours (**Figure 1A**). The inner diameters of the completed cuffs range from approximately 350 μm to 650 μm. The average irradiance produced by the silicone cuff-embedded LEDs was 6.4 ± 1.0 mW/mm². Optical transmittance through the vagus nerve was estimated using a one-dimensional modified Beer-Lambert law^38^ and optical properties of the rat sciatic nerve at 472 nm^39^. At a depth of 180 μm, the estimated diameter of the mouse vagus nerve, our model predicts 72% transmittance, resulting in an estimated irradiance of 4.61 mW/mm² at the far surface of the nerve (**Figure 1C**). The irradiance suggests adequate optical intensity to activate ChR2 throughout the entire nerve, based on published estimation of ChR2 activation threshold.^40, 41^

**Figure 1:**
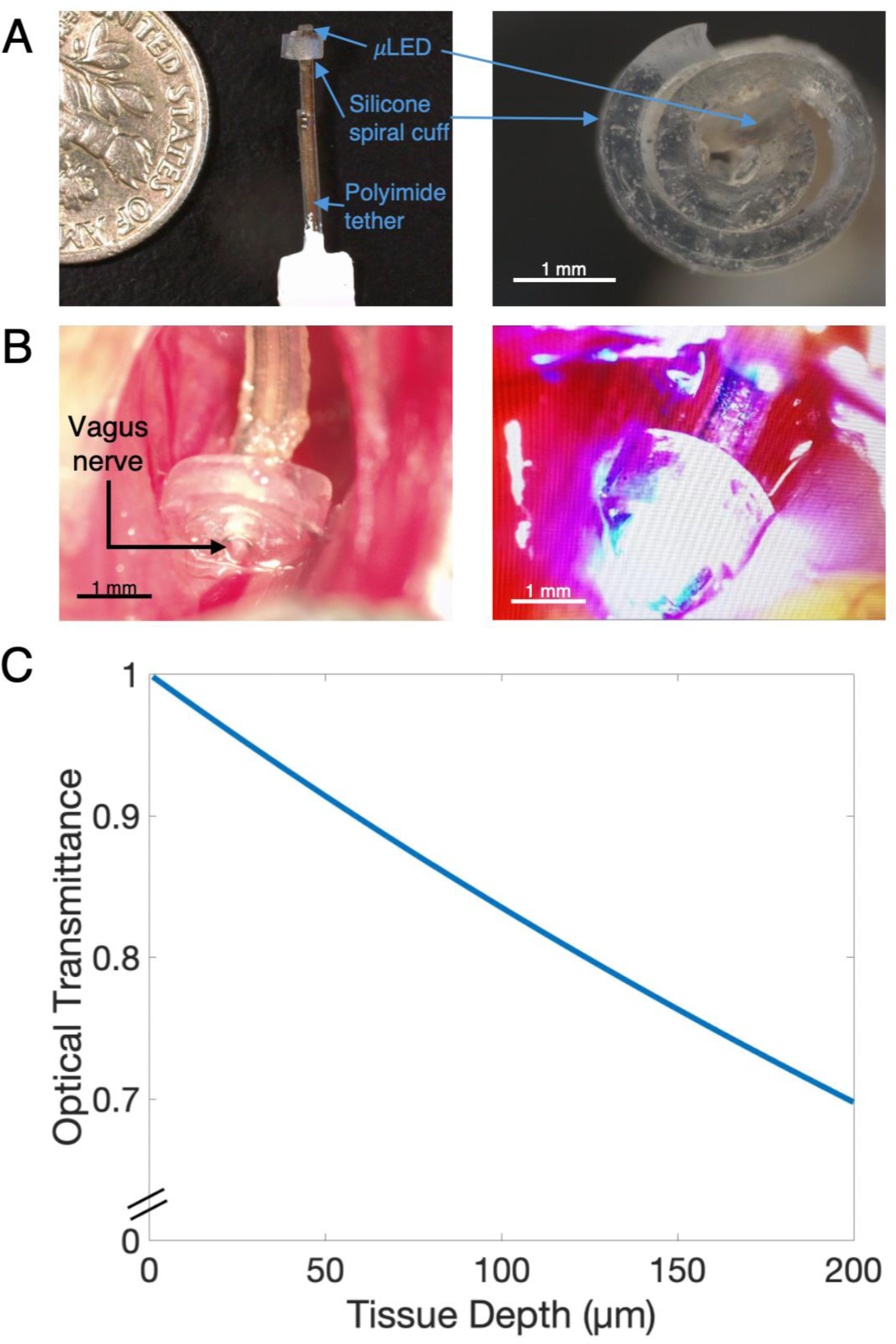
Silicone spiral nerve cuff with embedded μLED. **A:** Representative silicone spiral nerve cuff with integrated µLED probe **B:** Nerve cuff implanted on the left cervical vagus nerve of mice, shown in its ‘off’ (left) and ‘on’ (right) states. **C:** Theoretical optical transmittance of the device in nerve tissue as a function of tissue depth.

### Animals

The use of animals was approved by the Institutional Animal Care and Use Committee (IACUC) at the University of Colorado, Anschutz Medical Campus, with accreditation by the Association for Assessment and Accreditation of Laboratory Animal Care (AAALAC). All experiments were performed in accordance with IACUC regulations under an approved protocol. Transgenic mice (ChAT-cre (B6;129S6-*Chat^tm2(cre)Lowl^*/J), Vglut2-cre (B6J.129S6(FVB)-*Slc17a6^tm2^*^(cre)*Lowl*^/MwarJ), ChR2 (B6.Cg-*Gt(ROSA)26Sor^tm32(CAG-COP4*H134R/EYFP)Hze^*/J)) were obtained (Jackson Laboratory) and crossed to breed two experimental lines: ChAT-cre x ChR2 (referred to herein as ‘ChAT’), and Vglut2-cre x ChR2 (referred to as ‘Vglut2’). Cre^-^ non-expressing mice were used for ‘Sham’ stimulation. Adult mice of mixed sex between 15 and 30 weeks were used (*Supplementary Data Table S1*).

### Vagus Nerve Cuff Implantation

Anesthesia was induced with 5% isoflurane (Fluriso™) and maintained at 1–3% via nose cone. The mouse was placed in the supine position, and fur was removed from the ventral cervical region with Nair. A ventral cervical incision was made 2–3 mm from midline, and blunt dissection technique was used to expose the carotid sheath containing the carotid artery and vagus nerve. The vagus nerve was separated from the carotid sheath and surrounding tissue and implanted with a silicone spiral nerve cuff **(Figure 1B)**. Mice were euthanized upon completion of stimulation and blood draw. Mouse vitals were monitored throughout all procedures. Internal temperature was measured and maintained (RightTemp Jr ®; Kent Scientific) via rectal probe and a heating pad (37.5 °C) on the surgical plate. Other vitals (heart rate, pulse distension, breath rate, breath distension, pO2) were monitored and recorded via paw sensor (MouseOx Plus, Starr Life Sciences).

### Endotoxemia

Lipopolysaccharide (LPS) from *Escherichia coli* (O111:B4, Sigma Aldrich) was prepared in a 0.5% LPS, 0.9% saline solution and aliquoted and stored at -20 °C. Prior to use, aliquots were thawed and sonicated at room temperature for 30 minutes. Experimental LPS was delivered via intraperitoneal injection (3 mg/kg) 30 minutes post initialization of VNS protocol.

### Vagus Nerve Stimulation Parameters

Optical VNS was applied using 473nm light (2 Hz, 5 ms pulse width, 120 minutes) and controlled using nScope software.

### Serum Collection and Cytokine Measurement

Blood samples were collected (120 minutes after endotoxin injection, via cardiac puncture: 3CC syringe, 22G needle) and dispensed into clot activator tubes (CAT) (Greiner Bio-One GmbH). Samples were left to coagulate at room temperature for 30 minutes, then centrifuged (2000 x g, 20 minutes), and stored at -80°C until analysis. IL-1β, IL-6, IL-10, TNF-α and acute-phase protein, CRP, concentrations were assayed using a custom multiplex assay (U-PLEX Assay, Meso Scale Diagnostics) and a singleplex (R-PLEX CRP Assay, Meso Scale Diagnostics). All assays were run according to manufacturer protocols. Sample plates were quantified using the MESO QuickPLEX SQ120 instrument, and data was analyzed using MSD Workbench 4.0 Software.

### Fluorescence Microscopy

Bilateral excision of the cervical vagus nerves was performed post euthanasia. Tissues were fixed (4% paraformaldehyde, 30 minutes, phosphate buffered saline (PBS) wash) and mounted with media containing 40,6-diamidino-2-phenylindole (DAPI). Mounted tissue slides were stored at –20°C until analysis. Nerve samples were analyzed post-experimentally to image and verify fluorescently tagged ChR2 expression using a spinning disc confocal microscope (Marianas, 3i). Image processing was performed via FIJI software (*Supplementary Figure S3*).

### Statistical Analysis

Statistical outliers were identified using the mean absolute deviation method. Experiments that contained undue issues (i.e. deteriorating vitals; device malfunction) were identified as anecdotal outliers. Four data points appeared as both statistical and anecdotal outliers and were subsequently removed from the data set. Data is presented as mean ± SEM unless otherwise noted. Statistical analyses between groups were performed in MATLAB software using one-way ANOVA followed by the Least Significant Difference (LSD) post hoc test. P ≤ 0.05 was considered statistically significant.

## Results

Vagal stimulation was applied for two hours via implanted nerve cuff (**Figure 2A**) in acutely endotoxic (LPS-injected) mice in three experimental groups, with optogenetic ChR2 stimulation targeted to (1) efferent (ChAT) neuronal fibers, (2) afferent (Vglut2) fibers, and (3) a non-expressing Cre^−^ (Sham) control group. Blood was drawn via cardiac puncture for cytokine measurement 30 minutes following vagal stimulation (**Figure 2B**).

**Figure 2:**
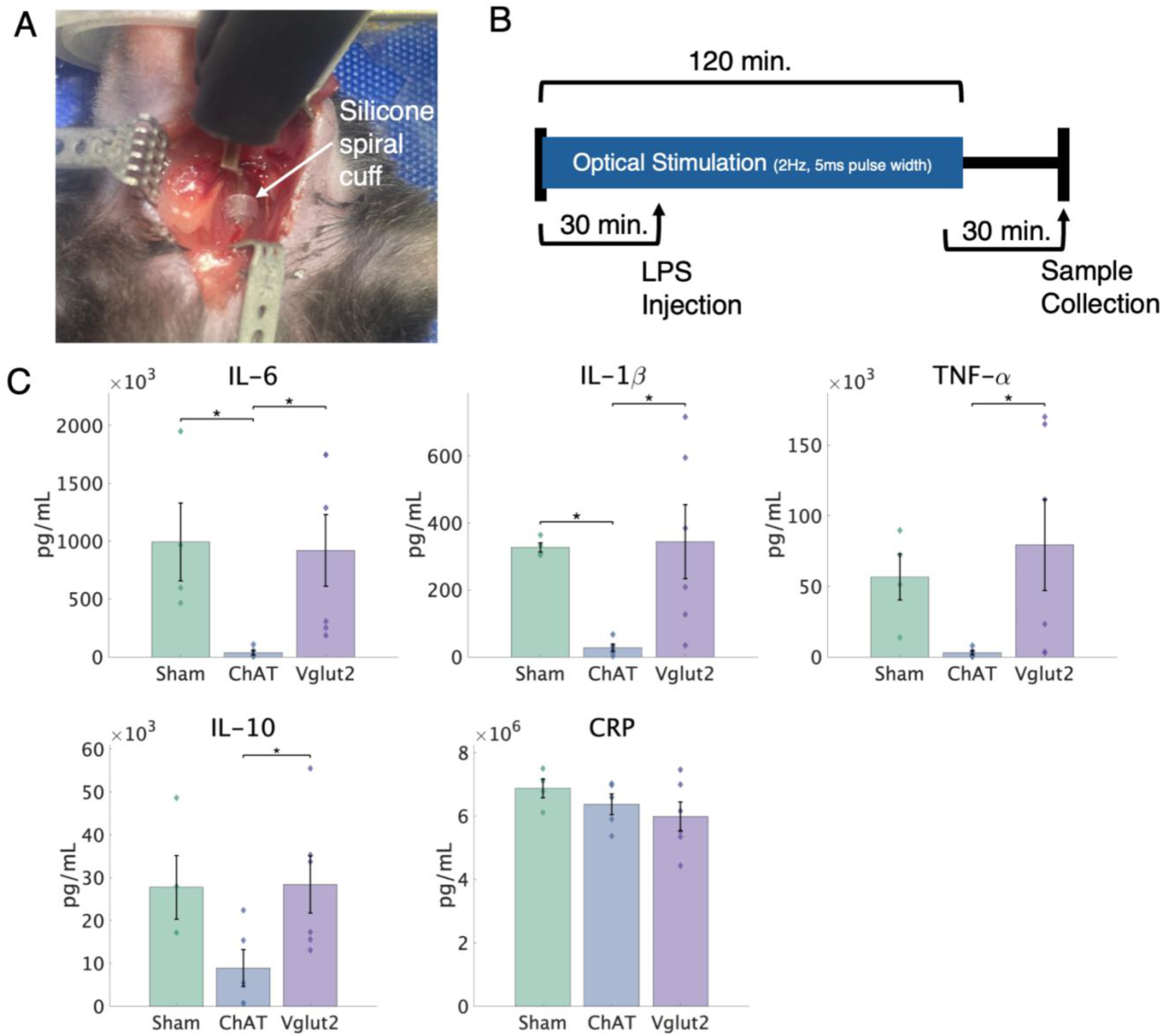
In vivo nerve cuff placement and experimental timeline for vagus nerve stimulation with resulting cytokine and CRP responses. **A:** Silicone spiral cuff implanted on the left cervical vagus nerve of an anesthetized experimental mouse. **B:** VNS protocol. All groups received VNS (2 Hz, 5 ms pulse width, 120 minutes), LPS (3 mg/kg via i.p. injection), and samples collected 120 minutes after LPS administration. **C:** Measurement of serum cytokine levels in LPS-challenged mice that received optogenetic VNS in selective axonal subsets with: ChAT-ChR2 expression in ChAT^+^ neurons (n=5), Vglut2-ChR2 expression in Vglut2^+^ neurons (n=6) and Cre^-^ Sham mice (n=4). Top: ChAT stimulation significantly reduced pro-inflammatory cytokines IL-6 and IL-1β relative to Sham controls (p=0.027 and p=0.026, respectively) and ChAT levels were reduced compared to Vglut2 stimulation in all three pro-inflammatory cytokines measured (IL-6: p=0.024, IL-1β: p=0.011, TNF-α: p=0.030). Bottom left: serum levels for anti-inflammatory cytokine IL-10 were reduced in the ChAT group compared to the Vglut2 group (p=0.033). Bottom right: serum levels for CRP were not statistically changed between groups. Mean ± SEM. * = p < 0.05.

Optogenetic VNS via ChAT neurons significantly reduced circulating serum concentration of pro-inflammatory cytokines IL-6 (p = 0.027), and IL-1β (p = 0.026) compared to the Sham stimulation group (**Figure 2C**). Surprisingly, ChAT stimulation did not significantlyreduce TNF-α levels relative to Sham (p = 0.15). With ChAT stimulation, IL-10 levels did not significantly differ from Sham (p = 0.058). Optogenetic stimulation of Vglut2 neurons did not significantly reduce the concentration of pro-inflammatory or anti-inflammatory cytokines when compared to Sham group: IL-6 (p = 0.85), IL-1β (p = 0.88), TNF-α (p = 0.51). IL-10 (p = 0.94). CRP levels did not significantly change from Sham with ChAT stimulation (p= 0.44) or Vglut2 stimulation (p=0.17).

However, the effect on pro-inflammatory cytokine levels did significantly differ between ChAT and Vglut2 stimulation groups: IL-6 (p = 0.024), IL-1β (p = 0.011), and TNF-α (p = 0.030) (**Figure 2C**). The anti-inflammatory cytokine IL-10 levels under ChAT stimulation were reduced relative to the Vglut2 group (p = 0.033).

In short, ChAT stimulation reduced serum cytokine concentrations compared with controls, whereas Vglut2 stimulation had no detectable effect on cytokine levels. ChAT and Vglut2 stimulation thus had markedly different effects on serum cytokines, with ChAT consistently reducing levels relative to Vglut2. This difference was observed for the pro-inflammatory cytokines IL-1β, IL-6, TNF-α, as well as the anti-inflammatory cytokine IL-10.

## Discussion

This study employed selective vagus nerve stimulation using optogenetic techniques to investigate the impact of distinct axonal subsets on inflammatory mediators in an acute model of systemic inflammation. Cholinergic (ChAT) and glutamatergic (Vglut2) vagal fibers were targeted separately to assess the relative contribution of these subsets to the attenuation of inflammatory cytokines. The study also sought to test whether the specific targeting of neuronal subsets with genetically defined activation yields robust modulation of inflammatory cytokines, demonstrating a potentially advantageous approach compared to electrical stimulation, which cannot achieve true cell-type specific stimulation.

Optogenetic VNS targeting cholinergic neurons significantly reduced IL-6 and IL-1β compared to sham, with a nonsignificant but concordant trend toward lower TNF-α levels, indicating a robust anti-inflammatory effect. These findings align with the established role of the CAP and prior work showing that activation of efferent neurons in the basal forebrain and vagus nerve decreases TNF-α and IL-6 levels in sepsis and endotoxemia models.^15, 20, 27, 42, 43^ ChAT stimulation produced 92-96% reductions in pro-inflammatory cytokine levels relative to sham, exceeding the 25-50% reduction typically reported in comparable electrical VNS studies.^27, 28^ While electrical stimulation favors larger fibers, optical stimulation can effectively recruit smaller cholinergic fibers of the CAP. Though variations in protocol across studies do exist, cell type specific recruitment with optogenetic stimulation is a likely contributor to the enhanced effect seen here, as ChAT optogenetic VNS selectively engages cholinergic efferent fibers and avoids the broad, nonspecific fiber activation inherent to electrical VNS that may engage pathways with opposing immunomodulatory actions. Nonetheless, these data support efferent CAP activation as a key mechanism for VNS-mediated pro-inflammatory cytokine suppression in acute systemic inflammation.

ChAT VNS did not significantly alter the anti-inflammatory cytokine IL-10, though mean levels were less than Sham. Prior efferent-targeted VNS studies report reduced or unchanged IL-10, while reductions in proinflammatory cytokines are more consistent.^15, 16, 42^ IL-10 may be more context-dependent and have different timescale dynamics. Measurement of cytokines at a greater range of timepoints will be needed in the future to better understand the impact of CAP on IL-10. CRP levels were unchanged in this experimental model. This is not unexpected given the delayed kinetics of CRP as an acute-phase protein, where peak levels may have occurred beyond the timeline of our experiment.^44–51^ Future experiments with extended timelines or alternative markers, such as serum amyloid P (a critical acute-phase reactant in murine models), may provide additional insight into the effects of VNS on downstream inflammatory processes.

Optogenetic stimulation of glutamatergic neurons did not significantly modulate inflammatory cytokines in the present experiments. This contrasts from reports that afferent electrical VNS modulates cytokine levels^20, 27, 52^, often through the engagement of non-cholinergic sympathetic pathways.^22, 53, 54^ Studies targeting specific afferent subpopulations (Vglut2^+^, TRPV1^+^, CALCA) show varied cytokine modulation across disease models.^19, 21, 43, 52^ Collectively, these findings indicate that afferent VNS may exert context-dependent effects that depend on the recruited subpopulation and inflammatory setting. It is also possible that afferent pathways impact inflammatory processes with different temporal dynamics compared to the efferent cholinergic pathway. Thus, disease specific studies with greater temporal characterization of inflammatory cytokines, as well as broader tissue and symptom-based assessment, will ultimately be needed to determine optimal axonal targets.

The lack of effect with Vglut2 neuronal activation, alongside the robust cytokine suppression observed with ChAT stimulation, underscores the importance of pathway specificity in VNS mediated inflammatory control. Efferent cholinergic and afferent glutamatergic fibers recruit distinct neuroimmune circuits, indicating that the inflammatory consequences of VNS depend critically on which vagal pathways are engaged.

A fundamental distinction between electrical and optogenetic VNS is specificity of neural activation. Electrical VNS recruits a heterogeneous mix of afferent and efferent fibers, engaging multiple physiological pathways and causing off-target effects. This nonselective recruitment may contribute to outcome variability and-limit the potential effect of isolated anti-inflammatory mechanisms, in contrast to the cell-type specificity achievable with genetically targeted-VNS. Optogenetic strategies therefore provide a powerful approach to dissect vagal circuits underlying inflammatory control and to design more precisely targeted neuromodulation therapeutics. The enhanced suppression of inflammatory cytokines observed with selective ChAT VNS relative to many studies have important implications for clinical translation, suggesting that improved disease treatment may be possible with these approaches.

## Conclusions

Optogenetic VNS targeting cholinergic (ChAT) neurons yielded significant reductions in IL-1β and IL-6 consistent with robust engagement of the cholinergic anti-inflammatory pathway. Stimulation of glutamatergic (Vglut2) neurons did not significantly alter inflammatory cytokine levels and, across all cytokines, responses to ChAT and Vglut2 VNS were significantly different, reinforcing that the inflammatory consequences of VNS are pathway specific and highly dependent on vagal subpopulation. These pathway-dependent effects highlight the value of optogenetic approaches for dissecting vagal circuits and suggest that selectively targeting efferent cholinergic fibers may be particularly effective for next generation neuromodulatory therapies for inflammatory disease. Future studies should include a broader time scale of cytokine measurements to better assess glutamatergic contribution and influence on IL-10 and acute phase proteins. Targeting additional vagal subpopulations beyond ChAT and Vglut2 neurons (for example TRPV1⁺ or CALCA⁺ populations) will help map the circuits underlying VNS-mediated inflammatory control and may reveal new therapeutic targets. Chronic optogenetic VNS studies in models of persistent inflammation are also needed to assess long-term impact while optimizing stimulation parameters to maximize therapeutic benefit.

