## Supplemental Information for "Attenuation of Inflammatory Cytokines by Selective Vagal Motor Stimulation via Silicone Spiral Nerve Cuff"

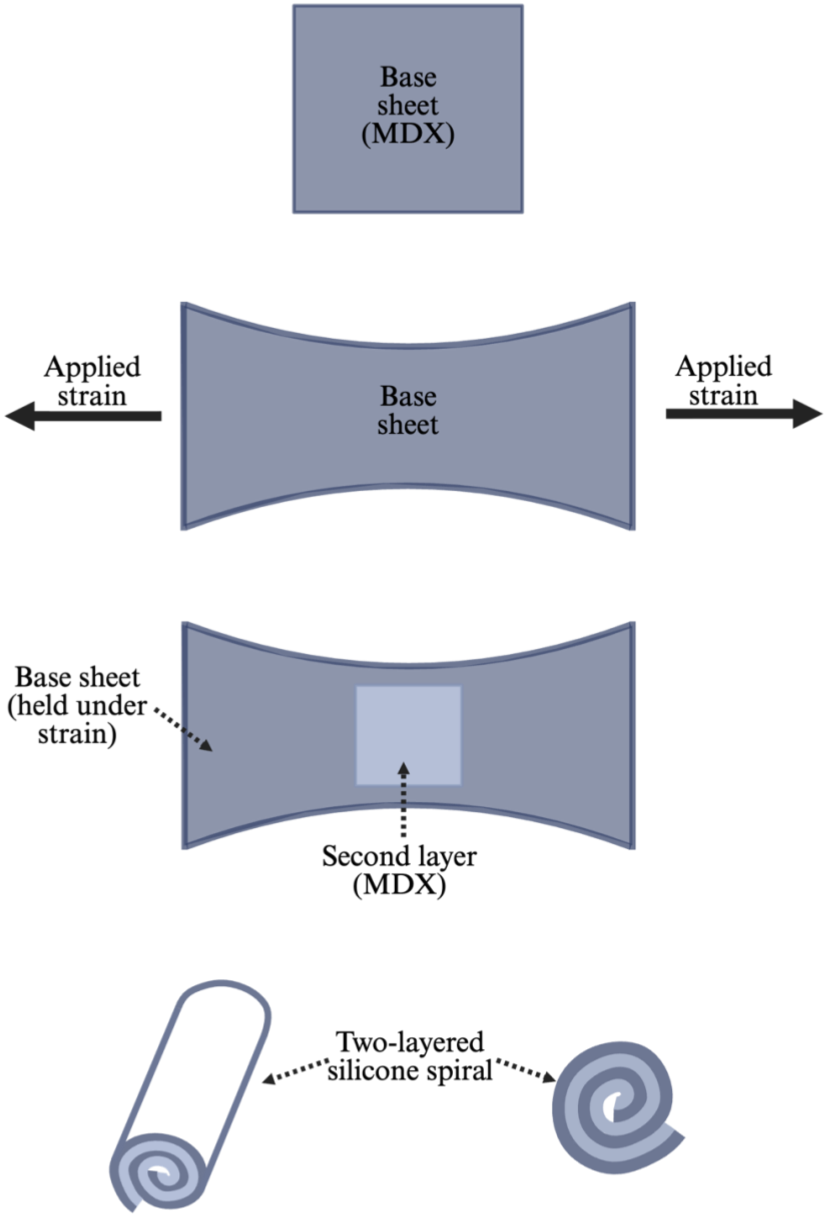


**Supplementary Figure S1:** Schematic of the silicone spiral nerve cuff manufacturing process. From top to bottom: silicone base sheet; strain applied to base sheet; second silicone layer (light blue square) cured on top of the strained base sheet; fully cured, two-layer, silicone spiral in oblique (left) and straight (right) views after demolding. Created in BioRender. Mirandette, K. (2026) https://BioRender.com/f8y8uj2.

| **Experimental Group** | Sham | ChAT | Vglut2 |
|---|---|---|---|
| **Number of Subjects (n)** | 4 | 5 | 6 |
| **Sex (M/F)** | 2/2 | 3/2 | 4/2 |
| **Age (min-max in weeks)** | 19-24 | 16-30 | 17-29 |
| **Transgenic Line**<br>*Genotype – (n)* | ChAT x ChR2<br>*WT/HZ – 1* | ChAT x ChR2<br>*HZ/HZ – 5* | Vglut2 x ChR2<br>*HZ/HZ – 2*<br>*HET/HET -3*<br>*Unknown – 1* |
| | Vglut2 x ChR2<br>*WT/HZ – 1*<br>*WT/HET – 1* | | |

**Supplementary Table S1:** Mouse cohort details. HZ = homozygous. HET = heterozygous. WT = wild type.

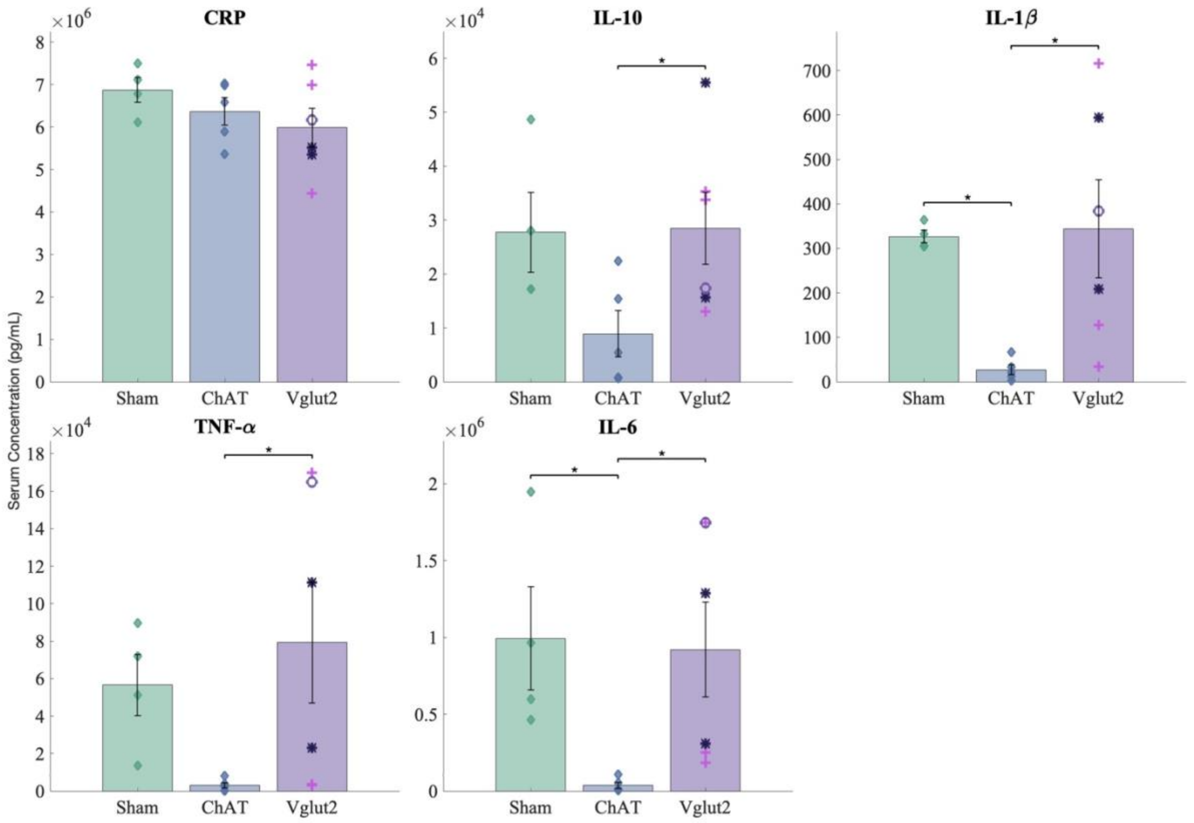


**Supplementary Figure S2**. Cytokine and CRP serum level results from inflammation experiment with marked Vglut2 genotypes. In the Vglut2 group, genotype is denoted for each subject data point with an overlayed shape. HZ/HZ = dark purple asterisk (✱). HET/HET = pink cross (✚). Unknown = purple circle (○). There is no significant difference in cytokine levels between the (HET/HET) and (HZ/HZ) groups; p > (0.5 for all analytes). Mean ± SEM. * = $p < 0.05$.

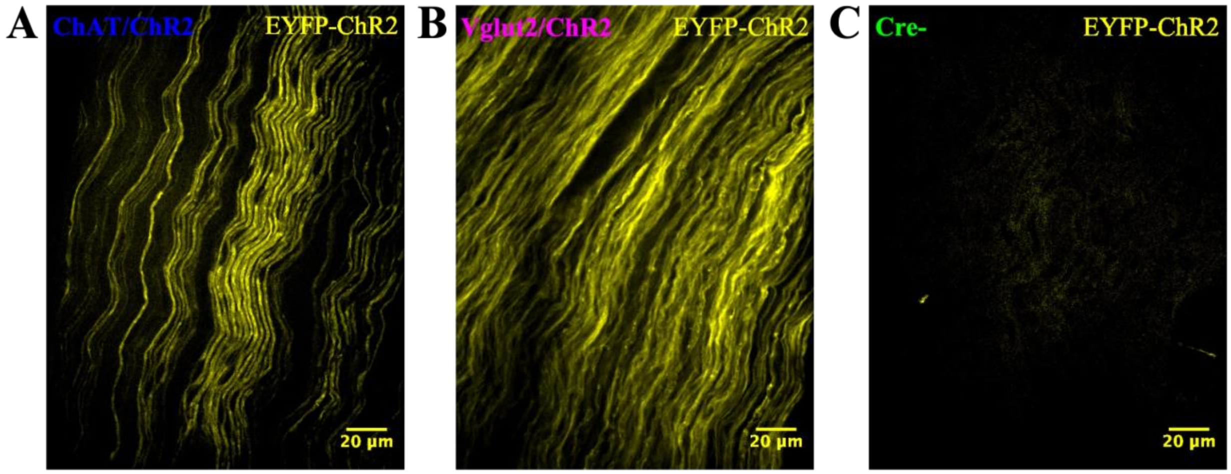


**Supplementary Figure S3:** Fluorescence microscopy of ChR2 expression in mouse vagus nerves. Tissue samples were analyzed via spinning disc confocal microscopy (excitation 515 nm, 40X/NA1.30 oil-immersion objective). ChR2 was tagged with Enhanced Yellow Fluorescent Protein (EYFP). **A:** Example positive ChR2 expression in a ChAT/ChR2 mouse. **B:** Example positive ChR2 expression in a Vglut2/ChR2 mouse. **C:** Example negative ChR2 expression in Cre- control mouse. Created in BioRender. Mirandette, K. (2026) https://BioRender.com/m632eh9.
